# Hyperactive somatostatin interneurons near amyloid plaque and cell-type-specific firing deficits in a mouse model of Alzheimer’s disease

**DOI:** 10.1101/2022.04.27.489759

**Authors:** Moustafa Algamal, Alyssa N. Russ, Morgan R. Miller, Steven S. Hou, Megi Maci, Leon P. Munting, Qiuchen Zhao, Dmitry Gerashchenko, Brian J. Bacskai, Ksenia V. Kastanenka

## Abstract

Alzheimer’s disease (AD) is characterized by synaptic loss and neuronal network dysfunction. These network deficits are mediated by early alterations in neuronal firing rates that coincide with amyloid plaque accumulation. Mounting evidence supports that inhibitory networks are impaired in AD, but the mechanisms driving these inhibitory deficits are poorly understood. Here we use in vivo multiphoton calcium imaging to determine the relationship between amyloid accumulation and the spontaneous activity of excitatory neurons and inhibitory interneurons in an APP/PS1 mouse model of Alzheimer’s disease. We show that somatostatin-expressing (SOM) interneurons are hyperactive, while parvalbumin-expressing interneurons are hypoactive in APP/PS1 mice. Only SOM interneuron hyperactivity correlated with proximity to amyloid plaque. These inhibitory deficits were accompanied by decreased excitatory neurons activity and decreased pairwise activity correlations in APP/PS1 mice. Our study identifies cell-specific interneuronal firing deficits driven by amyloid pathology in APP/PS1 mice and provides new insights for targeting inhibitory circuits in Alzheimer’s disease.

## Introduction

Alzheimer’s disease (AD) is the most common form of dementia and the most prevalent neurodegenerative disorder in the United States^1^. The primary pathological hallmarks of Alzheimer’s disease include deposits of extracellular amyloid plaques and intracellular tau tangles, and eventually neuronal death^2,3^. Notably, the accumulation of amyloid-beta peptides is associated with aberrant neuronal activity and oscillatory network alterations in AD patients^4,5^ and in animal models^6–9^. Particularly, deficits in cognition-linked brain rhythms such as gamma and slow waves are prevalent in AD patients^10,11^. These neuronal activity alterations appear at the early stages of AD and correlate with the severity of cognitive impairment in AD patients 12,13. Oscillatory brain rhythms are generated by complex firing patterns of individual neurons and interactions between neuronal populations across different brain regions^14–16^. Thus, there is an urgent need to understand the single-cell neuronal firing behavior in AD in order to restore the impaired oscillatory brain rhythms and improve cognitive function.

Several studies have assessed single-cell neuronal activity in amyloidosis mouse models^6,17,18^. However, since these reports did not discriminate between neuronal subtypes, the activity of excitatory cells was the predominant measure as they represent 80-90% of cortical neurons^19^. These reports have provided mixed findings of aberrant neuronal hyperactivity ^6,7^ and hypoactivity^6,20,21^. It is important to note that the hyperactivity phenotype has been observed in amyloidosis mouse models exhibiting a considerable number of seizures ^22,23^, and it remains unknown whether this phenotype occurs in amyloidosis models lacking the seizure phenotype. Answering this question is essential to understanding whether the hyperactivity phenotype is related to amyloid plaque accumulation or plaque unrelated factors. Thus, it remains unclear if network alterations in AD are due to increased or decreased excitability of excitatory cells.

In addition to excitatory cells, around 10-20% of cortical neurons are inhibitory interneurons that show a wide variety of morphological and electrophysiological properties^19,24–26^. Despite constituting a small percentage, interneurons play an essential role in cortical computations and the maintenance of spontaneous network oscillations^25^. Interneurons are classified based on their protein expression profiles into distinct subtypes including somatostatin (SOM)- and parvalbumin (PV)-expressing interneurons, and others^27^. SOM and PV interneurons shape excitatory activity by targeting different cellular compartments of pyramidal cells^25^.

The function of specific interneuronal cell types is impaired in AD^28,29^. Particularly, PV interneuron dysfunction received considerable attention because of its role in gamma oscillatory activity^28,30^. For instance, hippocampal PV interneurons are hypoactive in amyloidosis mouse models^30,31^. The firing activity of SOM interneurons in AD mouse models remains unexplored, but impairments in synaptic rewiring of hippocampal SOM interneurons was reported^29^. These cell-type-specific dysfunctions disrupt the well-regulated excitation and inhibition (E/I) balance resulting in broader, circuit-level dysfunction in AD^28^. Whether all interneuronal cell types are affected similarly in AD and whether their function is directly affected by amyloid plaques remains unclear.

During action potential firing, calcium ions (Ca^2+^) enter neurons through voltage gated calcium channels resulting in changes in intracellular Ca^2+^ concentration^32^. As result, calcium imaging has been widely used as a proxy of neuronal action potential firing^17,33,34^.

Here, we determined the effect of amyloid-beta accumulation on the spontaneous activity of three distinct neuronal populations within cortical layers 2/3. To this end, we monitored the activity of excitatory neurons and two major classes of inhibitory interneurons, PV and SOM, in anesthetized APP/PS1 mice and non-transgenic littermates (WT) using the genetically encoded calcium indicator GCaMP.

We show that SOM interneurons are hyperactive, while PV and excitatory neurons are hypoactive in APP/PS1 mice. Furthermore, SOM interneurons located near amyloid plaques are more active than those located farther away. Finally, pairwise neuronal correlations of excitatory neurons are decreased in APP/PS1 mice.

## Results

### SOM interneurons are hyperactive near amyloid plaque in APP/PS1 mice

The impact of AD on inhibitory interneuron cell-type-specific spontaneous activity is not well understood, with cortical SOM interneuron activity totally unexplored in vivo. Here, we performed in vivo multiphoton calcium imaging in anesthetized APP/PS1 (APP) mice and non-transgenic littermates (WT), to study cell-type-specific spontaneous neuronal activity.

We first monitored calcium transients in the somas of SOM interneurons using the genetically encoded calcium indicator jGCaMP7s (Figure1a). To target SOM interneurons selectively, we injected FLEX-jGCAMP7s into the somatosensory cortex of WT-SOM and APP-SOM mice (Figure 1a). SOM interneurons showed spontaneous calcium transients in WT-SOM (Figure 1b) and WT-APP (Figure 1c) mice. We then estimated the number of deconvolved calcium events as a proxy measure for neuronal action potential firing (Supplementary Figure 1, Movie 1). Our results show that SOM interneuron event rates were 2.7-fold higher in APP-SOM mice relative to WT-SOM mice (Figure 1d, 1e). The fraction of inactive SOM cells was not different between conditions (Figure 1f). Thus, SOM interneurons are hyperactive in APP/PS1 mice.

**Figure 1.**
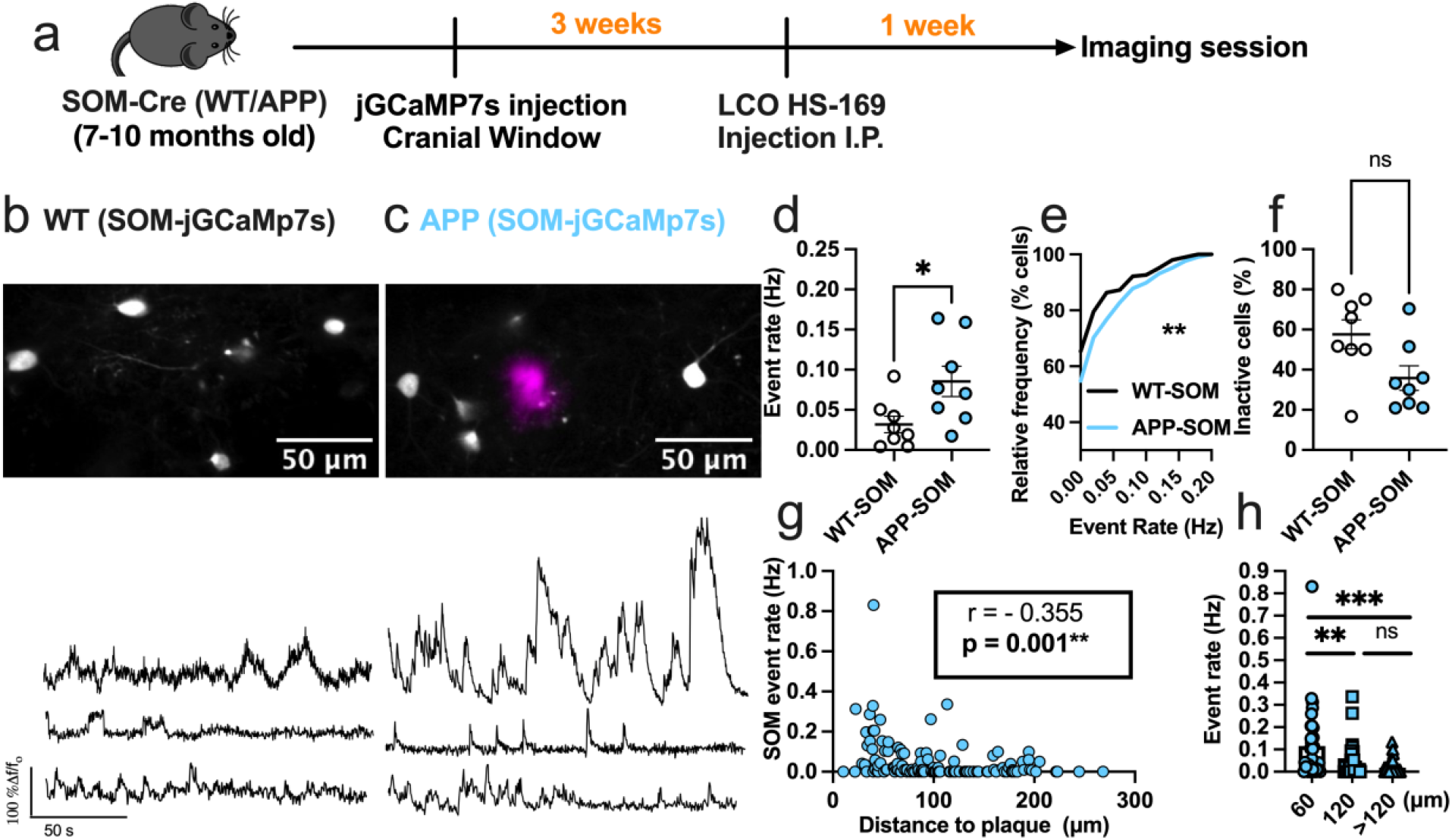
SOM interneurons are hyperactive in APP/PS1 mice relative to WT mice. **a**, Timeline of the experimental procedures. **b-c**, Top, in vivo two-photon fluorescence images of jGCaMP7s (grey) in SOM expressing interneurons in layer 2/3 of the somatosensory cortex from WT (**b**) and APP/PS1 (**c**) mice. Amyloid plaques were labeled with LCO-HS-169 (magenta); Scale bars, 50 μm. Bottom, representative normalized fluorescence traces of SOM interneurons activity from control (**b**) and APP/PS1 (**c**) mice. **d**, Mean neuronal activity rates as determined by counting the rate of deconvolved Ca+2 events (Mann–Whitney *U* = 11, *p*= 0.024, two-tailed, n= 8 WT-SOM mice; 8 APP-SOM mice). **e**, Cumulative frequency distribution of event rates in all imaged neurons (KS-test:D= 0.152, p= 0.0044, n= 210 WT-SOM neurons; 364 APP-SOM neurons). **f**, Quantification of the fraction of inactive cells in APP-SOM mice relative to WT_SOM (Mann–Whitney *U* = 14, *p*= 0.06, two-tailed, n= 8 WT-SOM mice; 8 APP-SOM mice). **g**, Correlations between event rates and the distance to amyloid plaque center in SOM interneurons (Spearman correlation: r(124) = -0.355, p <0.0001. **h**, Quantification of pooled event rates as a function of distance to amyloid plaque. Neurons were pooled into three groups depending on their distance to amyloid plaque: <60 µm, 60-120µm, and >120µm (Kruskal-Wallis test, H(2) = 18.41, p < 0.0001). Post hoc multiple comparisons were p= 0.0028 for SOM event rates <60µm vs. SOM <120µm and p= 0.0001 for SOM event rates <60µm vs. rates >120µm and not significant, p= 0.715, for SOM event rates <120µm vs. rates >120µm. Each solid circle in **d, f** represents an individual animal, while each circle in **g**-**h** represents an individual neuron. All error bars reflect the mean ± s.e.m. Asterisks denote statistically significant differences (**p<0.01, ****p<0.0001), while ‘ns’ denotes no significance p>0.05.

Studies examining indiscriminate neuronal populations have shown that neurons near amyloid plaques at distances less than 60 µm are more active than those farther away^6^. Therefore, to understand the relationship between amyloid plaques and the observed hyperactivity, we measured the correlation between SOM interneuron event rates and distance to amyloid plaques in APP/PS1 mice. We found a significant negative correlation between SOM interneuron activity and distance to amyloid plaque in APP/PS1 mice (Figure 1g). SOM interneurons near amyloid plaques (<60 µm) were more active compared to those located away from plaques (Figure 1h). Therefore, SOM interneuron hyperactivity is correlated with proximity to amyloid plaques. Postmortem immunohistochemical analysis of WT-SOM mice confirmed that 90 % of jGCaMP7s cells were positive for SOM (Supplementary Figure 2a, b), thus jGCaMP7s targeted SOM interneurons with high selectivity.

### PV interneurons are hypoactive in APP/PS1 mice

We then asked whether this hyperactivity can be generalized to other interneuron subtypes in APP/PS1 mice. PV interneurons have markedly different electrophysiological properties relative to SOM interneurons as they can reach very high firing rates^35^. Hence, we next used similar experimental procedures (Figure 1a) and targeted jGCaMP7s expression to PV interneurons in WT-PV-Cre (WT-PV) (Figure 2b) and APP-PV-Cre (APP-PV) mice (Figure 2c).

**Figure 2.**
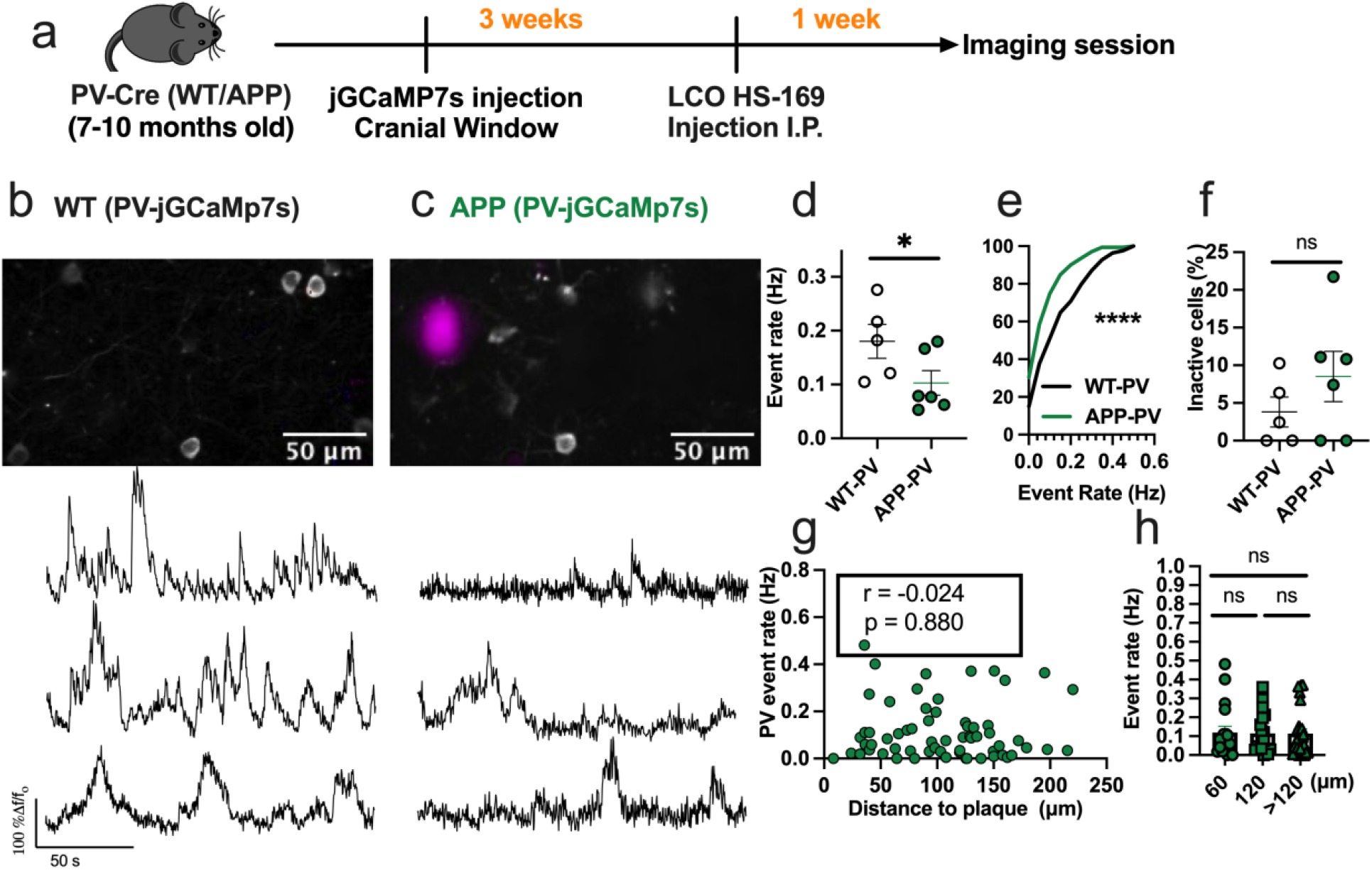
PV interneurons are hypoactive in APP/PS1 mice relative to WT mice. **a**, Timeline of the experimental procedures. **b-c**, Top, in vivo two-photon fluorescence images of jGCaMP7s in PV expressing interneurons (grey) in layer 2/3 of the somatosensory cortex from WT (**b**) and APP/PS1 (**c**) mice. Amyloid plaques were labeled with LCO-HS-169 (magenta); Scale bars, 50 μm. Bottom, representative normalized fluorescence traces from control(**b**) and APP/PS1 (**c**) mice. **d**, Mean event rates (Mann–Whitney *U* = 4, *p*= 0.051, two-tailed, n= 5 WT-PV mice ; 6 APP-PV mice). **e**, Cumulative frequency distribution of event rates in all imaged neurons (KS-test:D= 0.268, p< 0.0001, n= 254 WT-PV neurons; 172 APP-PV neurons). **f**, Quantification of the fraction of inactive cells (Mann–Whitney *U* = 4, *p*= 0.28, two-tailed) in imaged WT-PV and APP-PV (n= 5 WT mice; 6 APP/PS1 mice). **g**, Correlations between event rates and the distance to amyloid plaque center in PV interneurons (Spearman correlation: r(63) = -0.024, p = 0.88), and **h**, quantification of pooled PV event rates as a function of amyloid plaque distance (Kruskal-Wallis test, H(2) = 0.46, p < 0.79). Each solid circle in **d**-**f** represents an individual animal, while each circle in **g**-**h** represents an individual neuron. All error bars reflect the mean ± s.e.m. Asterisks denote statistically significant differences (*p<0.05, ****p<0.0001), while ‘ns’ denotes no significance p>0.05.

Mean event rates were 5.6-fold higher in WT-PV interneurons (Figure 2d, Movie 2) relative to WT-SOM interneurons (Figure 1d) (mean ± s.e.m.: 0.18±0.03 for PV interneurons, and 0.032±0.01 for SOM interneurons). Contrary to our findings in SOM interneurons, PV interneurons showed a 1.8-fold decrease in event rates in APP-PV mice relative to WT-PV mice (Figure 2d, 2e). The fraction of inactive PV cells was comparable across conditions (Figure 2f). PV interneuron activity did not correlate with proximity to amyloid plaques (Figure 2g, h).

Therefore, PV interneurons are hypoactive in APP/PS1 mice irrespective of their proximity to amyloid plaques. These results suggest a cell-type-specific disruption in cortical interneuron activity of APP/PS1 mice.

Postmortem immunohistochemical analysis of PV-Cre mice confirmed that 86 % of jGCaMP7s cells were positive for PV (Supplementary Figure 2c,d).

### Excitatory neurons are hypoactive in APP/PS1 mice

The disrupted interneuronal activity of APP/PS1 mice prompted us to investigate whether this deficit is related to E/I imbalance. SOM and PV interneurons form inhibitory synaptic contacts onto excitatory neurons, therefore a dominant hypoactivity in either or both interneuron subtypes can potentially result in hyperactivity of excitatory cells. Because of the numerous reports of hyperactive cells in APP/PS1 mice, we hypothesized that excitatory cells are hyperactive as result of a dominant hypoactivity in PV interneurons. To test this hypothesis, we imaged spontaneous activity of layer 2/3 cortical excitatory neurons in WT (WT-EX) and APP/PS1 (APP-EX) mice. We targeted excitatory neurons using a human CaMKIIα promoter to drive the expression of GCaMP6s (Figure 3a-c). These recordings were performed on a combination of SOM-and PV-Cre mice to ensure comparability to interneuron recordings (see methods). In contrast to our hypothesis, excitatory neurons of APP-EX mice showed 1.6-fold lower event rates (Figure 3d, Movie 3) relative to WT-EX mice, with an overall lower cumulative distribution of event rates across all recorded cells (Figure 3e). Mean event rates of excitatory cells were stable over two to three months (Supplementary Figure 3) and did not depend on sex or mouse model (Supplementary Figure 4). No change in the fractions of inactive excitatory cells (Figure 3f) was observed. Excitatory neuron activity did not depend on amyloid plaque distance (Figure 3g, 3h, Movie 4). Therefore, cortical excitatory neurons are hypoactive in APP/PS1 mice irrespective of their proximity to amyloid plaques.

**Figure 3.**
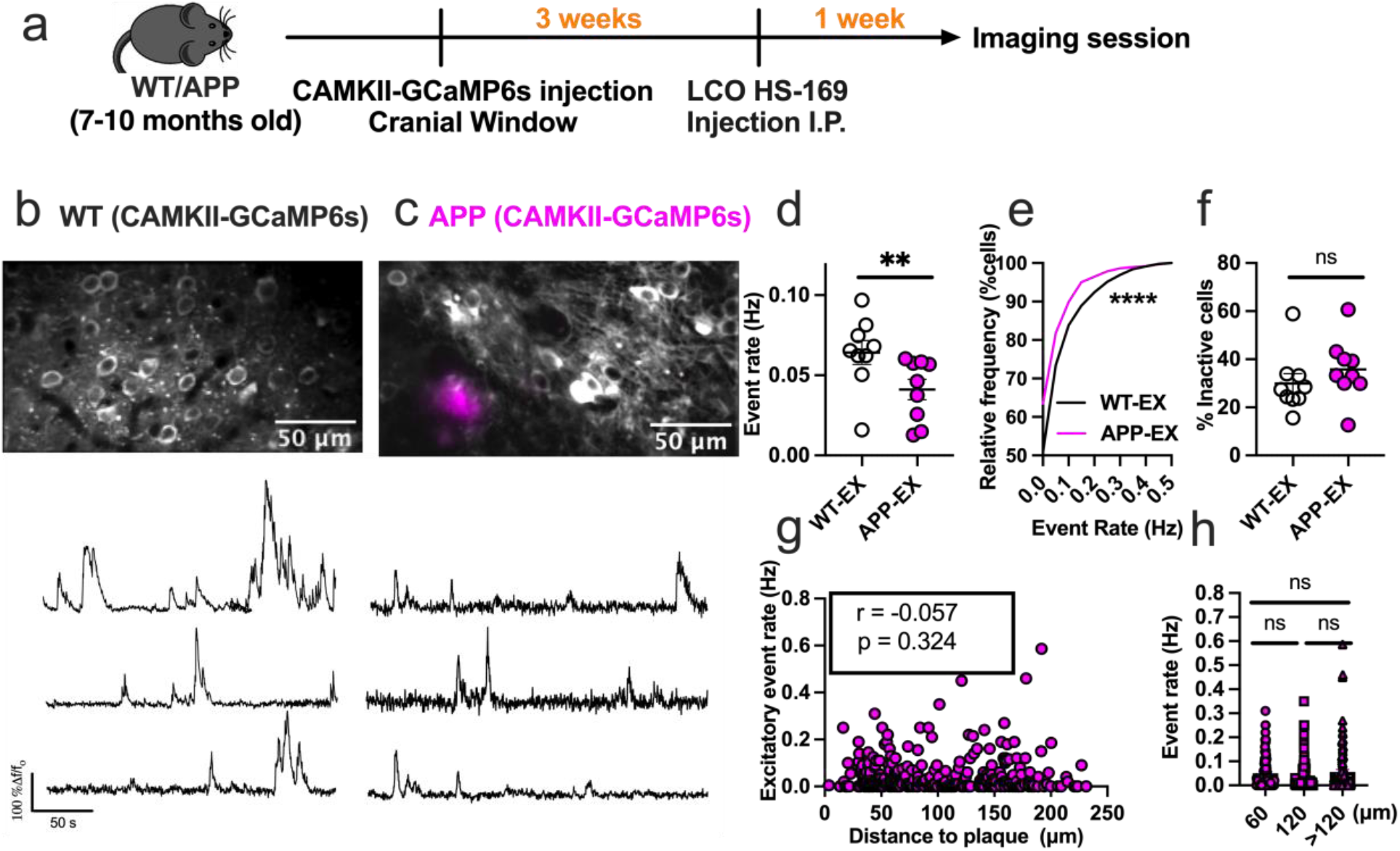
Excitatory neurons are hypoactive in APP/PS1 mice. **a**, Timeline of the experimental procedures. **b-c**, Top, in vivo two-photon fluorescence images of GCaMP6s in CaMKII expressing (excitatory) neurons (grey) in layer 2/3 of the somatosensory cortex from non-transgenic controls (**b**) and APP/PS1 (**c**) mice. Amyloid plaques were labeled with LCO-HS-169 (magenta); Scale bars, 50 μm. Bottom, representative normalized fluorescence traces from control (**b**) and APP/PS1 (**c**) mice. **d**, Mean neuronal activity rates as determined by counting the rate of deconvolved Ca+2 events (Mann–Whitney *U* = 11, *p*= 0.0078, two-tailed). **e**, Cumulative frequency distribution of event rates in all imaged neurons (KS-test:D= 0.129, p< 0.0001, n= 1412 WT-EX neurons; 1603 neuron APP-EX neurons). **f**, Quantification of the fraction of inactive cells (Mann–Whitney *U* = 23, *p*= 0.136, two-tailed) in imaged WT-EX and APP-EX (n= 9 WT-EX mice; 9 APP-EX mice).**g**, Correlations between event rates and the distance to amyloid plaque center in excitatory neurons (Spearman correlation: r(298) = -0.057, p = 0.324), and **h**, quantification of pooled excitatory neurons event rates as a function of amyloid plaque distance (Kruskal-Wallis test, H(2) = 0.46, p < 0.79). Each solid circle in **d**-**f** represents an individual animal, while each circle in **g**-**h** represents an individual neuron. All error bars reflect the mean ± s.e.m. Asterisks denote statistically significant differences (*p<0.05, ****p<0.0001), while ‘ns’ denotes no significance p>0.05.

### APP/PS1 mice show decreased pairwise excitatory neuronal correlations

To evaluate the impact of amyloid accumulation on firing synchrony of different cell types, we calculated the Pearson correlation values (ρ) for all possible pairwise combinations of active neurons. Mean correlation values were significantly lower in APP-EX relative to WT-EX mice (Figure 4a). It is established that neuronal synchrony decreases as the distance between neurons increases^36^. Therefore, we determined the relationship between correlation values and pairwise distance between neurons (Figure 4b). As expected, correlation values decreased as a function of increased inter-neuronal distance, a phenomenon that seemed to be similar in WT-EX and APP-EX mice (Figure 4b). Since correlations are known to increase when neuronal event rates increase^37^, we also examined the relationship between mean correlations of a given neuron and its event rate. Excitatory neuronal ρ values increased as function of neuronal event rates (Figure 4c). These observations did not significantly differ in WT-EX and APP-EX mice (Figure 4c), suggesting that the decreased correlation values in APP-EX mice are related to decreased event rates and not synchrony. We calculated significantly fewer pairwise correlations for interneurons relative to excitatory cells because only a few cells could be recorded in each imaging frame. Nevertheless, there was no detectable change in correlation values for SOM and PV interneurons in APP/PS1 mice relative to WT (Figure 4d, 4e).

**Figure 4.**
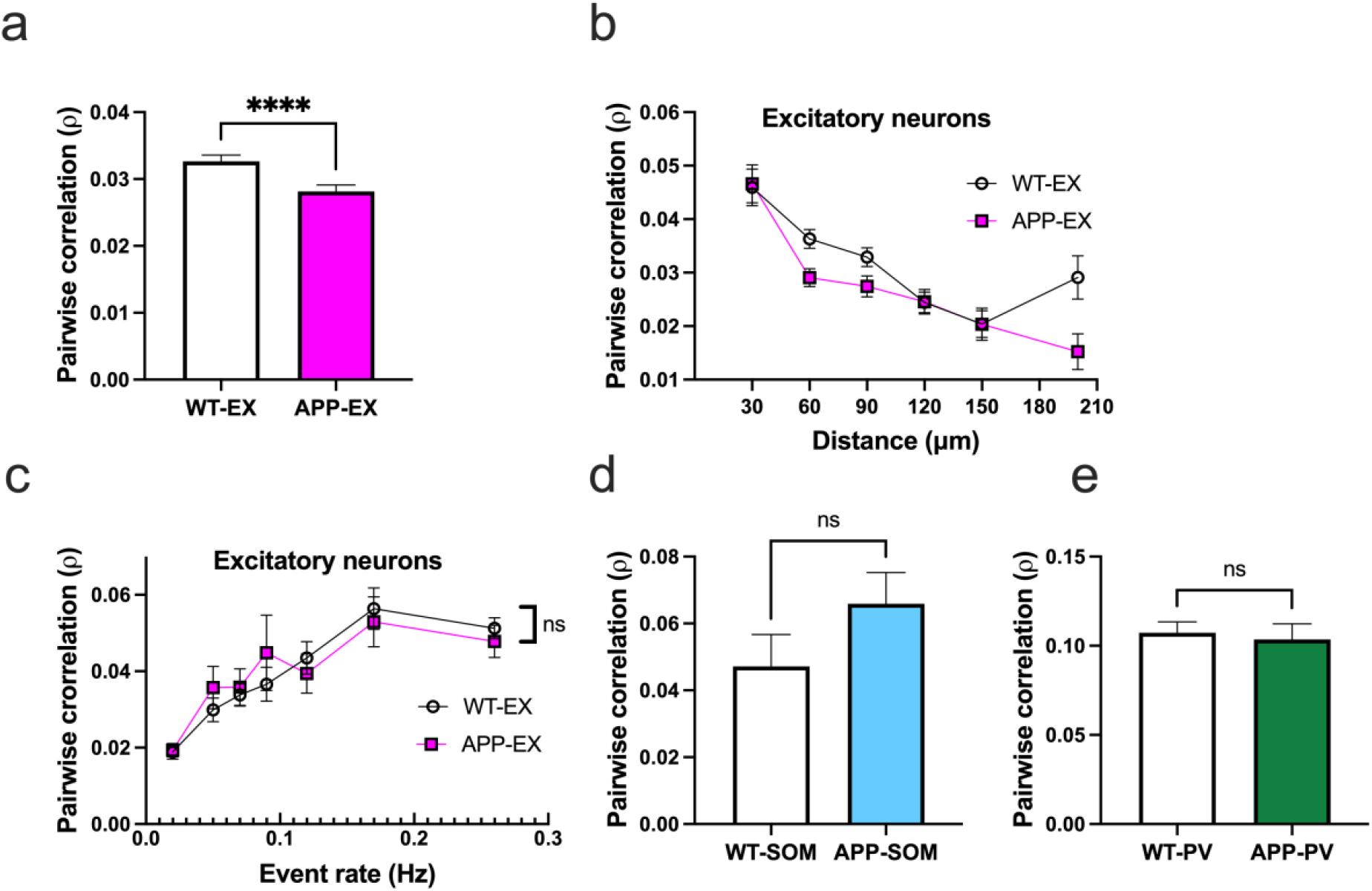
Decreased pairwise correlations in excitatory neurons of APP/PS1 mice. **a**, Average pairwise Pearson correlation values for excitatory neurons in APP/PS1 and WT mice (Mann–Whitney *U* = 3012, n = 8791 WT-EX pairwise-correlations, n = 8409 APP-EX pairwise-correlations, *p*<0.0001, two-tailed). **b**, Relationship between pooled pairwise Pearson correlation values and pairwise interneuronal distance (Two-Way ANOVA, main distance effect F_(5, 17193)_ = 21.78, p = <0.0001, main genotype effect F_(1, 17193)_ = 7.066, p = 0.008, main genotype and distance interaction F_(5, 17193)_ = 1.72, p = 0.128). The x axis values represent the maximum limit of the pooled pairwise distances. **c**, Relationship between correlation values and event rates after pooling data according to geometric means of neuronal event rates (Two-Way ANOVA, main event rate effect F_(6,2211)_ = 29.07, p = <0.0001, main genotype effect F_(1, 2211)_ = 0.74, p = 0.427, main genotype and event rate interaction F_(6, 2211)_ =0.55, p =0.77). **d-e**, Average pairwise Pearson correlation values for SOM interneurons (**d**) (Mann–Whitney *U* = 254487, n = 163 WT-SOM pairwise-correlations, n = 337 APP-SOM pairwise-correlations, *p*= 0.193, two-tailed) and PV interneurons (**e**) (Mann–Whitney *U* = 32424, n = 392 WT-PV pairwise-correlations, n = 166 APP-PV pairwise-correlations, *p*= 0.95, two-tailed) in APP/PS1 and WT mice. All error bars reflect the mean ± s.e.m.

## Discussion

Altered neuronal network activity in AD has been observed in several cerebral amyloidosis mouse models and AD patients^6,7,12,13,17,30,38,39^. Recent reports suggest that inhibitory circuits are particularly vulnerable, resulting in E/I imbalance and circuit-level dysfunction in AD^30,40–42^. However, the exact nature of the disrupted inhibitory activity is not yet fully understood. We report cell-type-specific interneuron dysfunction in APP/PS1 mice. SOM interneurons were hyperactive while PV interneurons were hypoactive. Additionally, the changes in SOM interneuron activity, but not PV activity, correlated with proximity to amyloid plaques. These inhibitory deficits were accompanied by a decrease in excitatory neuronal activity, thus providing a detailed picture of the nature of E/I imbalance in the APP/PS1 mouse model.

Our finding of decreased excitatory neuronal activity in APP/PS1 mice may appear to conflict with reports of hyperactivity in amyloidosis models^6,17,43^. Several factors could contribute to this discrepancy, including the population of neurons being investigated, animal model strain, and the data analysis methodology. Firstly, most studies reporting hyperactive cells did not distinguish among different neuronal cell types. It is possible that most of the previously reported hyperactive cells were, in fact, SOM interneurons. Furthermore, in agreement with our PV and excitatory cells findings, the initial reports of the hyperactivity phenotype also reported an increased fraction of hypoactive cells^6,44^. Secondly, we used APP/PS1 mice on C57BL/6;C3H background, which do not exhibit seizures^45^, while other amyloidosis models such as APP/PS1 mice on C57BL/6J background^7,17^ do exhibit seizures^45^. Since seizures could be accompanied by increased excitability of pyramidal cells^22,46^, higher seizure incidence could result in a dominant hyperactivity phenotype in an amyloidosis mouse model. On the other hand, the decrease in spontaneous activity of PV interneurons reported in this study is in agreement with previous reports linking PV interneuron deficits to gamma rhythm disruption in AD^7,28,30,31,47^.

Mounting evidence from in vivo and in vitro studies has shown that circuit activity is homeostatically regulated to maintain neuronal firing rates constrained to a particular functional limit^48,49^. We provide evidence for the failure of homeostatic mechanisms to maintain appropriate neuronal firing in APP/PS1 mice. Hyperactive SOM interneurons are the most likely driver of this dyshomeostasis since they supply aberrant inhibition to layer 2/3 pyramidal cells, thus preventing firing in excitatory cells^27,50,51^. The hyperactivity of SOM interneurons correlates with proximity to amyloid plaques. Thus, we propose a mechanism by which amyloid plaque-driven hyperactivity in SOM interneurons leads to decreased firing of pyramidal cells. SOM interneurons were also reported to strongly inhibit PV interneurons in cortical layers 2/3 during visual processing^52^. SOM interneuron hyperactivity could thus suppress PV interneuron firing in APP/PS1 mice. Indeed, we saw hypoactivity within PV interneurons.

Several in vitro studies have shown that the application of amyloid-beta oligomers induces neuronal hyperexcitability through several mechanisms that involve blocking glutamate uptake leading to glutamate spillover^41^, reduction of endocannabinoid-mediated disinhibition^53^, and increasing neuronal resting membrane potential^22^. A question that emerges from the current study is how amyloid beta-induced hyperactivity can be observed in SOM interneurons only and not in PV or excitatory cells. To answer this intriguing question, we adopt a network model in which two sources can perturb each neuronal cell-type activity: nearby amyloid plaque and excitatory and inhibitory network inputs. For instance, despite being conflicted by nearby amyloid plaque, excitatory neurons are hypoactive in APP/PS1 mice due to the hyperactive SOM inhibitory inputs. These opposite perturbations of excitatory cells can also explain the lack of correlation between excitatory cell activity and the distance to amyloid plaque.

Our results have important implications for future studies targeting the inhibitory circuit as a treatment strategy for AD. While inhibitory circuit deficits are prevalent in AD, it is not clear whether a therapeutic strategy that broadly restores inhibitory tone, using GABA agonists or antagonists, would benefit early-stage AD patients. Our results suggest that a more targeted approach to restoring inhibitory tone will be needed to overcome these network deficits since interneuron cell types were differentially affected by AD pathology. Future studies using approaches to control neuronal activity, such as optogenetics, should aim to decrease SOM or increase PV interneurons activity. Indeed, several studies have targeted PV interneurons activation using optogenetic and pharmacological approaches to restore oscillatory gamma activity and improve cognitive functions in preclinical animal models^28,31,47,54,55^.

Our study assumes that the Ca^+2^ indicator (GCaMP) concentration distribution across cells in different models is similar. A violation of this assumption could result in inaccurate estimation of event rates due to changes in indicator sensitivity and potentially cellular Ca^+2^ buffering capacity 56. Because there is no accurate way to estimate the intracellular GCaMP concentration, developing ratiometric indicators with comparable Ca^+2^ kinetics and sensitivity to the latest GCaMP series will be needed. We opted to perform our experiments under low isoflurane anesthesia to reduce experimental variability^57^ and compare our findings to previous reports of altered single-cell activity in AD^6,17^. However, anesthesia is known to reduce neuronal activity and presents a limitation to our current findings^58^. Therefore, future experiments should include reproducing these findings in awake behaving animals.

Together, these findings show an association between amyloid pathology and interneuron-related circuit deficits in an Alzheimer’s disease mouse model. These results should guide future therapeutic approaches targeting circuit dysfunction at the early stages of Alzheimer’s disease.

## Data availability

All data needed to assess the conclusions of this paper are present in the main text or the Supplementary Information. Data is available from the corresponding author upon reasonable request.

## Author contributions

Conception and design of study (M.A., K.V.K., B.J.B.), mouse surgery (M.A., A.N.R.), in vivo imaging and maintenance of two-photon setup (M.A., S.S.H.), data analysis (M.A. with help from M.M, L.M), coding and data interpretation (M.A.), immunohistochemistry (M.R.M), manuscript preparation (M.A. with help from A.N.R., B.J.B., K.V.K.), securing funding (K.V.K, B.J.B, D.G.) and project supervision (B.J.B, K.V.K).

## Methods

### Animals

All experimental procedures were approved by MGH Institutional Animal Care and Use Committee (IACUC). We crossed homozygote APPswe/PSdE9 transgenic mice overexpressing the amyloid precursor protein with the Swedish mutation and deltaE9 mutation in presenilin 1 (APP/PS1) (Stock #034829, C57BL/6;C3H genetic background, The Jackson Laboratory)^59^ with homozygote SOM-IRES-Cre knock-in mice (Stock #013044, The Jackson Laboratory)^60^ or homozygote PV-Cre mice (Stock # 008069, The Jackson Laboratory)^61^ to generate APP-SOM and APP-PV mice, respectively. Age-matched non-transgenic littermates were used as controls (WT-SOM and WT-PV). Experiments were performed in 8–11-month-old animals including males and females. Prior to any surgical procedures, animals of the same sex were housed in up to 4 animals/cage. Individual housing was maintained after surgical manipulation. All animals had ad libitum access to water and food and were maintained on a 12/12-hour day/night cycle in a pathogen-free environment.

### Cranial window surgery and viral transduction of genetically encoded Ca^2+^ indicator

Mice were initially anesthetized with 5% isoflurane inhalation in O2 and maintained on 1.5% isoflurane during the surgery. Throughout the procedure, mice were placed on a heating pad to keep the body temperature at ∼37.5 °C and their eyes were protected with an ophthalmic ointment. A 5 mm craniotomy was drilled over the right somatosensory cortex using sterile technique. The dura was kept intact and wet with saline.

To express the ultrasensitive genetically encoded fluorescent calcium indicators in cortical inhibitory interneurons, we injected FLEX-jGCaMP7s (Addgene# 104491-AAV1)^40^ in SOM-CRE (4× 10^12^ genome copies/ml) and PV-CRE (1 × 10^12^ genome copies/ml) WT and APP mice (n=5-8 per group) at ∼300 μm below the surface in the somatosensory cortex (2 sites, at 1.5- and 2.5-mm posterior to bregma and 2 mm lateral to the midline suture). We then used adeno-associated virus AAV1 carrying the construct GCaMP6s under the CaMKIIα promoter (Addgene# 107790-AAV9, 1 × 10^12^ genome copies/ml)^41^ to target excitatory cells of WT and APP mice (n=9 per group). To ensure our excitatory neurons experiments are comparable to our interneurons experiments, CaMKIIα-GCaMP6s injections were targeted to a combination of PV-CRE (n=5 per group) and SOM-CRE (n=4 per group) WT and APP mice All the injections were delivered using a Hamilton syringe and Pump 11 Elite microsyringe pump (Harvard Apparatus) at a rate of 200nl/min. Each injection consisted of ∼1 μl of viral construct diluted in phosphate-buffered saline containing 0.01% Pluronic F-68. After injecting the desired volume, the Hamilton syringe was left at the injection site for 5 min to prevent backflow of the viral solution. A round glass coverslip (5 mm diameter) was placed over the right somatosensory cortex using a mix of dental cement and cyanoacrylate. Following each surgery, mice were housed individually and received analgesia (buprenorphine and acetaminophen) for 3 days postoperatively.

### In vivo multiphoton calcium imaging

Imaging was performed under light isoflurane anesthesia to reduce experimental variability^12^. Mice were initially sedated with 5% isoflurane in room air using the SomnoSuite®Low-Flow Anesthesia System (Kent Scientific). Mice were then imaged under light anesthesia and low air-flow rates (1% isoflurane and ∼40 mL/min air flow for a 30 g mouse). A heating pad was used to keep the animal’s body temperature at 37.5 °C, and ophthalmic ointment was used to protect the eyes. Animals were kept for at least 60 minutes under light isoflurane anesthesia before imaging. Mice typically showed an absence of tail-pinch reflex and a respiration rate of 80-120 breaths per minute during the imaging session.

A Fluoview FV1000MPE multiphoton microscope (Olympus) with a mode-locked MaiTai Ti:sapphire laser (Spectra-Physics) set to 900 nm was used for two-photon imaging. Spontaneous Ca^2+^ fluorescence signals from the somatosensory cortex were collected at 5 Hz through a 25x, 1.05 numerical aperture water immersion objective (Olympus) at 2x digital zoom. The Fluoview program was used to control the scanning and image acquisition (Olympus). Multiple fields of view (160 x100 µm, 1 pixel per µm) were imaged per mouse, and each field of view was recorded for at least 100 seconds.

### Calcium imaging data analysis

We used Suite2P^62^ and a custom-written MATLAB program to analyze spontaneous calcium events as follows. Because of the slower timescale of calcium indicator responses relative to action potential-related voltage changes, deconvolution of the calcium signal to the best estimate of spiking activity is required^32,33^. Several calcium deconvolution algorithms have been developed to address this non-trivial issue^63–66^. We chose to use a non-negative deconvolution algorithm^66,67^ to estimate the timing and the number of deconvolved calcium events in our recordings (Movie 1, Supplementary Figure 1) because it outperformed all other algorithms in a dataset of mutual calcium imaging and ground-truth electrophysiology recordings^68^.

Recordings were collected as Olympus Image Format (OIF) files, converted to Tiff files using ImageJ, then imported to Suite2P for automated registration, somatic regions of interest (ROI) detection, and neuropil contamination estimation. We eliminated the neuropil contribution to emphasize neuronal soma signals. All images were aligned using the Suite2p rigid registration function that computes the shifts between each frame and a reference image using the phase correlation method. ROIs were automatically detected from the mean image for each recording using Suite2p anatomic detection methods. All automated ROIs were inspected manually, and missing ROIs were added. Fluorescence values were extracted from each ROI for each frame, and the mean for each cell was computed. In addition, neuropil contamination was estimated from at least 100 pixels surrounding each somatic ROI using Suite2p. This gave us two vectors of fluorescence values for the soma and the neuropil. Data were then analyzed using a custom-written MATLAB program that was adopted from prior calcium imaging studies ^34^ as described in the following paragraphs.

The neuropil vector was weighed a factor of 0.7^33^. The weighted neuropil vector was subtracted from the somatic vector to produce a corrected vector of fluorescence values. The normalized ΔF/F trace was calculated according to the equation (Fi-F0)/F0), where Fi is the frame index, and F0 is the fluorescence baseline. To measure baseline F0 fluorescence in each cell, we employed a moving average baselining function like prior calcium imaging investigations^34^. The baselining function applies a sliding window of 250 frames (50 seconds) and quartile cut-off values ranging from the bottom tenth percentile to the median, depending on how active the neuron was. We then counted the total number of deconvolved events from the generated ΔF/F traces to estimate neuronal event rates using a robust non-negative deconvolution approach^67^ (https://github.com/zhoupc/OASIS_MATLAB). The deconvolution was employed using a FOOPSI Autoregressive model#1 (AR1) with the event size constrained to be 2.5 times the noise levels. GCaMP6/7s decay time was set at 1.25s to estimate the ‘γ’ parameter, while sparsity penalty parameter (λ) was set to 0.

To calculate the correlation coefficients, deconvolved calcium events from each neuron were binarized and corrected for pixel timing using linear interpolation^69^. Correlation coefficients were then computed as the Pearson correlation coefficient (ρ) for all possible pairings of active neurons (event rate higher than 0.01 Hz) using MATLAB ‘corr’ function. All possible ρ values were pooled per experimental group to estimate the degree of synchrony. Distances between individual neurons represent the Euclidean distance between the ROI centroids. To estimate the effect of distance to each amyloid plaque on neuronal activity, we chose fields of view containing several neuronal cells and a single amyloid plaque in the same plane. Additionally, we made sure that no additional amyloid plaques were located within 100 µm in the x,y,z planes of the imaging field of view. Distances were then calculated as the Euclidean distances between the amyloid plaque center and the neuronal ROI centroid.

### Immunohistochemistry

After the acquisition of multiphoton images, SOM-Cre and PV-Cre mice injected with FLEX-jGCaMP7s were euthanized using CO2 inhalation. Animals were then transcardially perfused, their brains collected and kept in 4% paraformaldehyde (PFA)/ phosphate buffered saline (PBS) solution overnight. The following day, brains were washed in PBS then transferred to a cryopreserving solution (30% sucrose in PBS). 40 µm thick coronal slices of mouse brains were subjected to antigen retrieval in citrate buffer. The sections were subsequently permeabilized with Triton X-100, blocked with normal goat serum (NGS), and incubated overnight at 4°C with the following primary antibodies; PV (mouse monoclonal anti-PV, 1:500; Sigma P3088), SOM (rat polyclonal anti-SOM, 1:50; MAB354), or GFP (chicken polyclonal anti-GFP, 1:500, A10262). Tissue sections were then washed and incubated with their respective secondary antibodies (1:500) for 1 hour at room temperature. The slides were then mounted with Vectashield antifade mounting medium (Vector Laboratories). A Fluoview FV3000 confocal microscope (Olympus) with a 40x objective was used to image PV and SOM-stained sections. Quantification of SOM and PV positive cells was done manually.

### Statistics

The appropriate statistical test was chosen after the assessment of normal distribution using the Shapiro-Wilk normality test. In most cases, a non-parametric test was used because normal distribution could not be assumed. For comparison involving two experimental groups, we used either a parametric two-sided student t-test or a non-parametric Mann-Whitney t-test as detailed in the figure captions. Comparisons between three groups were assessed using Kruskal– Wallis one-way analysis of variance followed by a Dunn’s test to correct for multiple comparisons. Two-Way ANOVA was used to assess the relationship between inter-neuronal distances and pairwise neuronal activity correlations. Spearman correlation was used to assess the relationship between amyloid plaque distance and event rates. A p value of less than 0.05 was considered statistically significant.

