## Supplementary Figures for "Hyperactive somatostatin interneurons near amyloid plaque and cell-type-specific firing deficits in a mouse model of Alzheimer’s disease": Supplementary_MA2.docx

**for**

**Supplementary Figures**

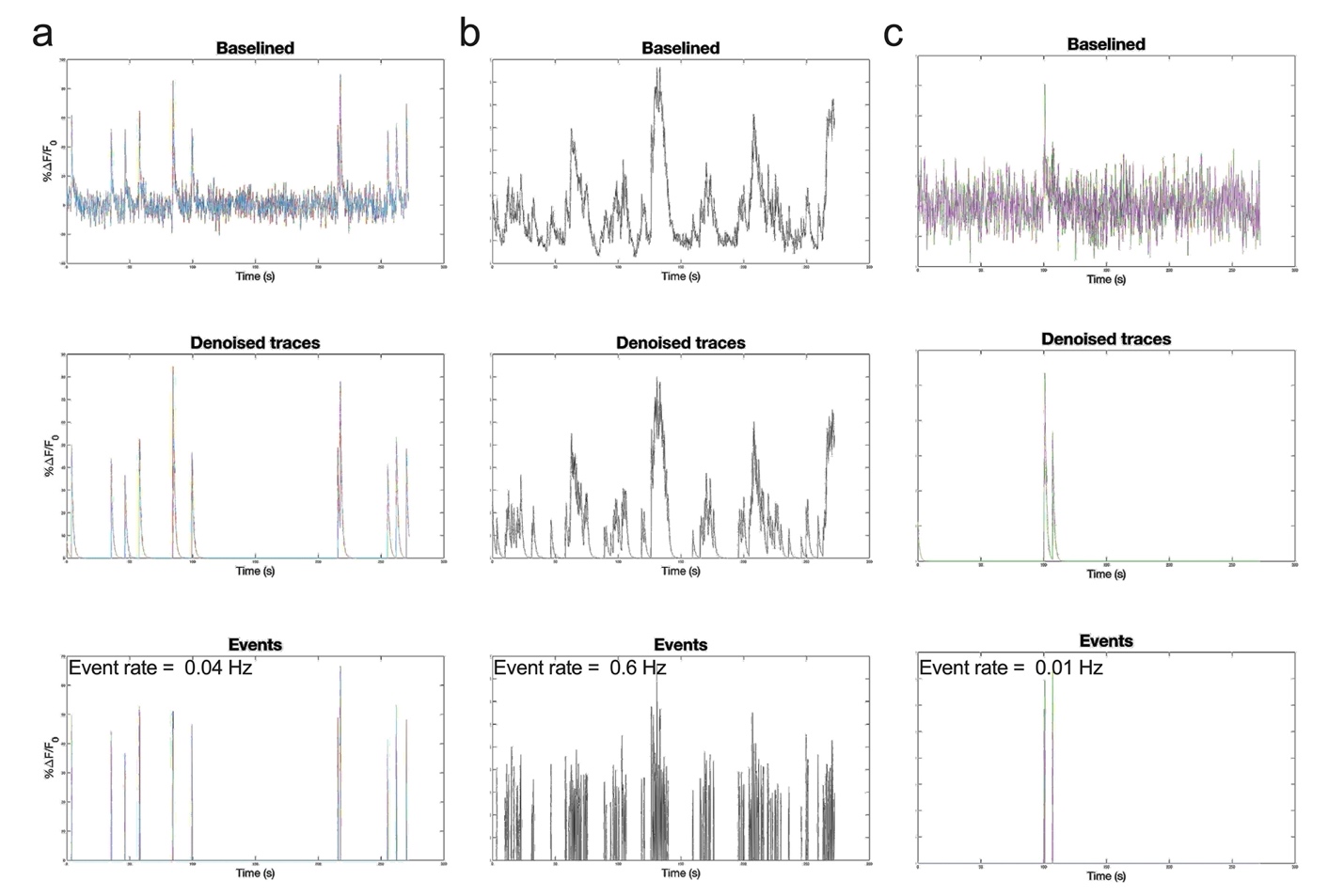

**Supplementary Fig. 1. Examples of deconvoluted calcium events in SOM interneurons**. Top, Baselined raw calcium traces from three SOM cells (**a**, **b** and **c**). See also a recording of the same cells in Movie 1. Middle, Denoised calcium traces from the same cells depicted in the top panels. Bottom, estimation of deconvolved calcium event rates after thresholding spikes to two and half times the standard deviation of the noise. We ran OASIS using the following MATLAB implementation:

[c_oasis, s_oasis, options] = deconvolveCa(y, 'ar1', g, 'foopsi', 'lambda', lambda, 'smin', smin). Lambda, the sparsity regularization parameter, was set to 0 while smin was set to -2.5.

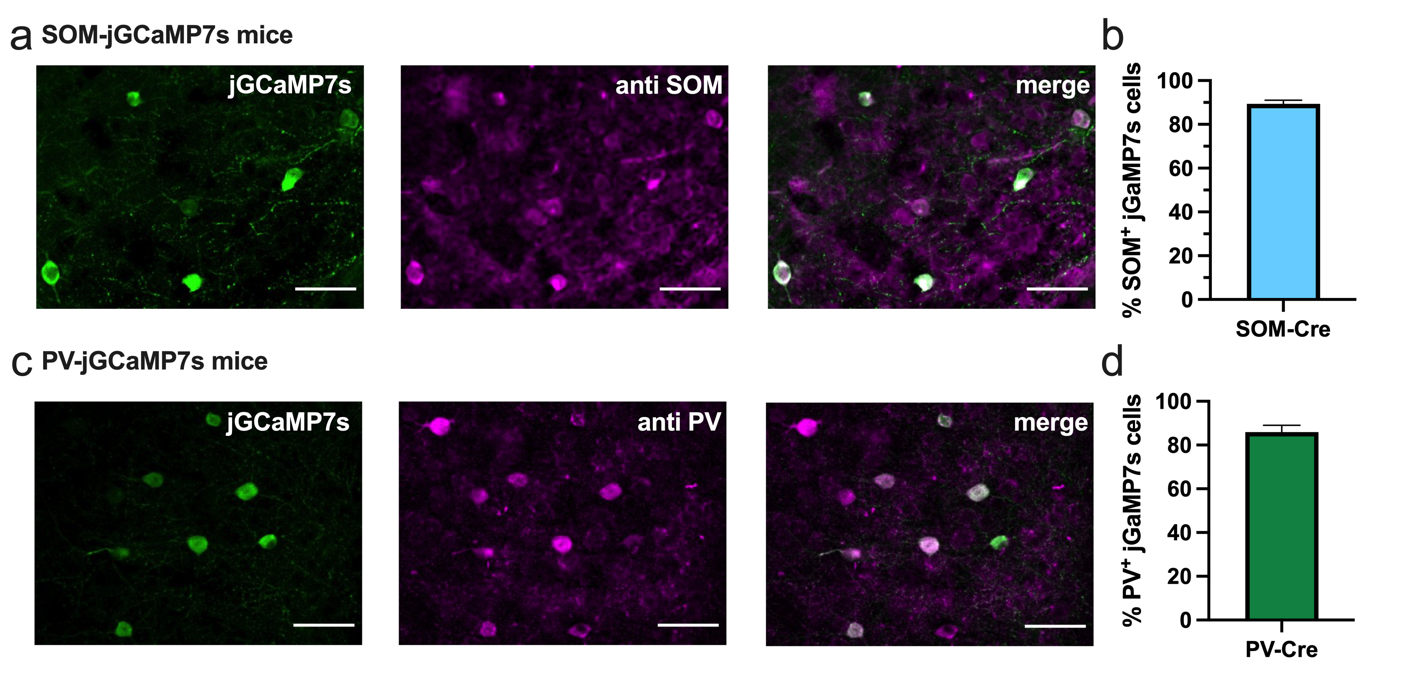

**Supplementary Fig. 2. Expression of jGCaMP7s in PV-Cre and SOM-Cre mice.**

Confocal images of a brain slice from a SOM-Cre mouse injected with FLEX-jGCaMP7s showing green jGCaMP7s (anti-GFP staining), SOM (anti-SOM staining) in magenta, and the merged image (**a**). Quantification of SOM positive cells show that 90 ± 1.6% of jGCaMP7s cells are SOM positive in SOM-cre mice (**b**). Confocal images of a brain slice from a PV-Cre mouse injected with FLEX-jGCaMP7s showing green jGCaMP7s (anti-GFP staining), PV (anti-PV staining) in magenta, and the merged image (**c**). Quantification of PV positive cells show that 86 ± 3.2% of jGCaMP7s cells are PV positive in PV-Cre mice (**d**). All error bars reflect the mean ± s.e.m.

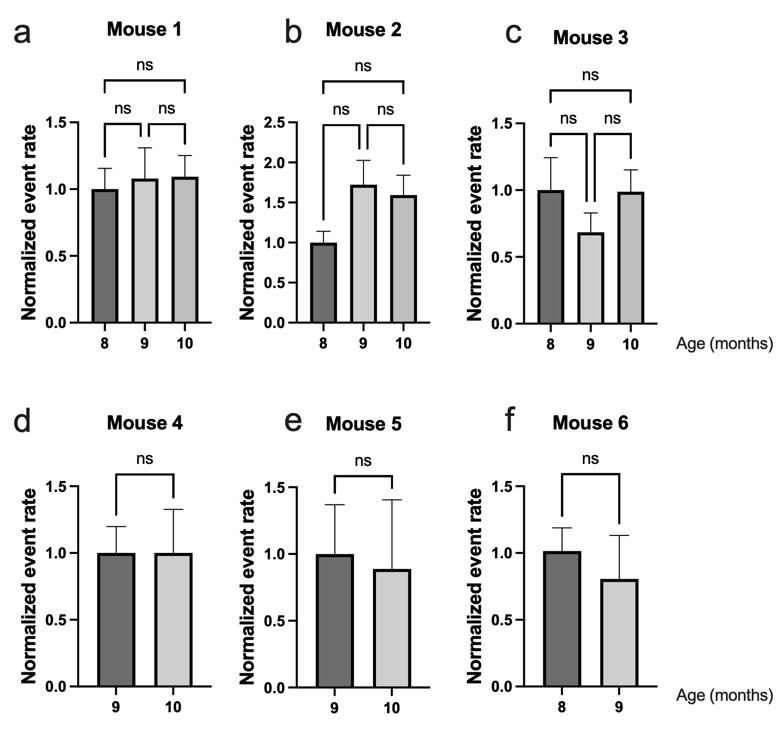

**Supplementary Fig. 3. Stability of neuronal event rates with age.** We monitored the activity of excitatory neurons in APP mice over multiple imaging sessions to examine the impact of aging on mean Ca+2 event rates. All event rates were normalized to the mean event rate during the first imaging session. No significant changes were observed in event rates in APP mice receiving multiple imaging sessions at 8, 9, or 10 months of age. Panels a-c were analyzed using One-way ANOVA followed by Dun’s test to correct for multiple comparisons, while panels e-f were analyzed using a t-test. Each bar represents the mean event rates per mouse, and all error bars reflect the mean ± s.e.m. ‘ns’ denotes no significance p>0.05.

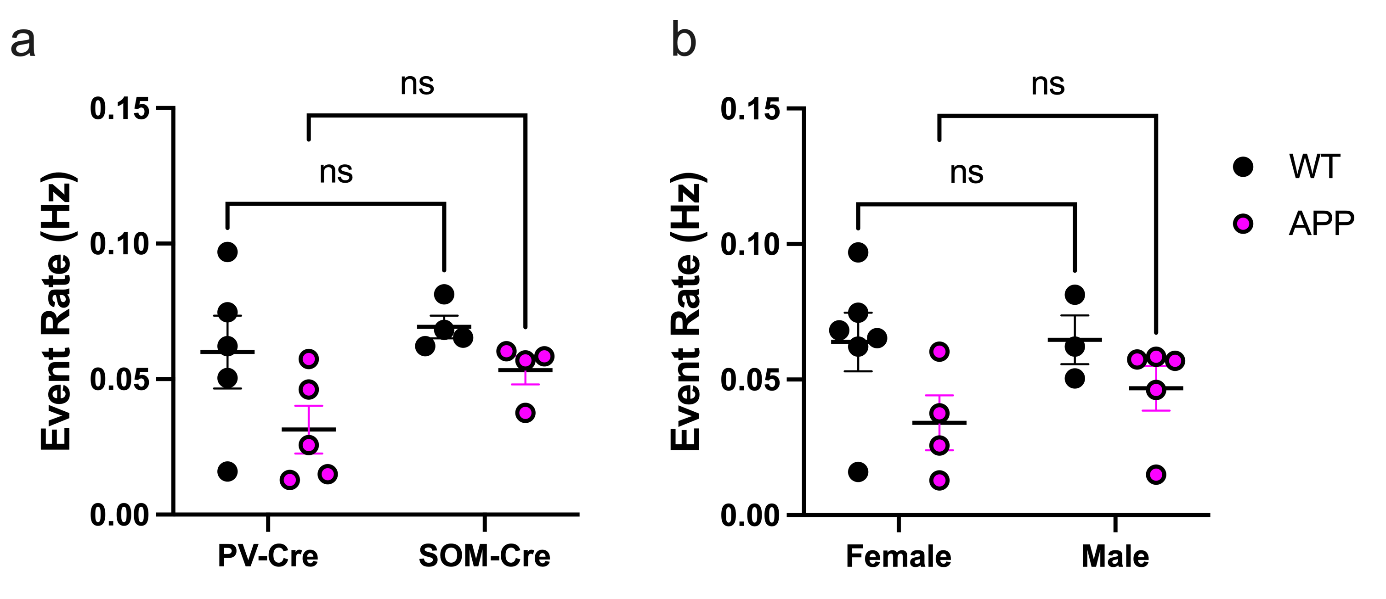

**Supplementary Fig. 4. Effect of mouse model and sex on excitatory cell event rates**. a, Mean event rates for excitatory cells from PV-Cre and SOM-Cre WT and APP mice (Two-Way ANOVA, main PV-Cre vs. SOM-Cre effect F_(1, 14)_ = 0.4, p = 0.54, main WT vs. APP effect F_(1, 14)_ = 5.07, p = 0.041, main interaction F_(1, 14)_ = 0.31, p = 0.58 ). b, Mean event rates for excitatory cells in male and female WT and APP mice (Two-Way ANOVA, main male vs. female effect F_(1, 14)_ = 2.6, p = 0.18, main WT vs. APP effect F_(1, 14)_ = 5.4, p = 0.035, main interaction F_(1, 14)_ = 0.44, p = 0.52 ). All error bars reflect the mean ± s.e.m.
